# An open-source application for applying rapid transient perturbations using a split-belt treadmill

**DOI:** 10.64898/2026.07.21.739794

**Authors:** Kiichi F. Ash, Courtney M. Butowicz, Brad D. Hendershot, Pawel R. Golyski

## Abstract

**Background:** The use of specialized perturbation systems has become an increasingly popular approach for investigating walking stability. By accelerating or decelerating one belt, researchers can induce slip- and trip-like perturbations in a controlled laboratory setting. However, many existing studies rely on specialized perturbation systems that require expertise in device-specific software and handling of the equipment, limiting the accessibility of perturbation-based gait research to laboratories with access to such equipment. To address this limitation, we developed an open-source method capable of inducing slip- and trip-like perturbations using a standard split-belt treadmill. Here, we 1) describe the hardware and software components of the system, 2) validate the application’s accuracy and precision, and 3) characterize the stability demands imparted by the perturbations with spatial stability measurements. Measured perturbation onset delay and duration were compared to the desired onset timing and programmed duration in addition to step length, step width, minimum mediolateral margin of stability, and sagittal-plane whole-body angular momentum range during the perturbed and recovery steps.

**Results:** Five participants with traumatic unilateral transtibial limb loss experienced perturbations consisting of brief, rapid increases or decreases in unilateral treadmill velocity, eliciting a “slip” or “trip”. The mean (standard deviation) onset delay was 183.3 (9.7) ms, or 24.18% (1.91%) of stance duration. Mean perturbation duration was 239.90 (7.5) ms, 18.14% longer than the intended duration. The perturbations produced measurable changes in gait stability, such as increased step length during the perturbed step and step width during the subsequent recovery step in addition to increased minimum mediolateral margin of stability and sagittal whole body angular momentum.

**Conclusion:** This open-source method successfully induced instability in individuals with impaired balance, demonstrating its feasibility as an accessible alternative to specialized perturbation systems. Future work will focus on refining both the software and hardware components to further improve timing, accuracy, and consistency.

## 1. Background

Losses of stability during walking are a significant problem, with >35% of older adults (1,2), > 34% of patients with neurological diseases (3), and >50% of individuals with lower limb loss falling at least once a year (4,5). In addition to injury (6,7), falls can precipitate a fear of falling, resulting in activity avoidance (4). Unexpected perturbations, such as trips and slips, are among the leading causes of falls during walking (7–9).

A mechanistic understanding of perturbation responses is crucial for developing rehabilitation interventions to reduce fall risk. Experiments in this context must be performed in controlled, safe, and data rich environments. Multiple methods have been used to impart unexpected mechanical perturbations in laboratory settings, including obstacles (10–12), robotic systems (13,14), and split-belt treadmills (15). Of these methods, split-belts treadmills are appealing since they leverage hardware already present in many research labs. However, implementing belt-speed perturbations on a split-belt treadmill is largely limited to specific high-performance devices that require expertise in device-specific software (15).

Here, this study details an open-source method that allows for rapid transient changes in belt speed to be applied at specified times during the gait cycle using a standard split-belt treadmill. This approach uses a low-cost data acquisition board to trigger short duration accelerations/decelerations of individual belts at a user-specified percentage of stance phase, as determined using vertical ground reaction forces. The objectives in this work are to: 1) describe the hardware and software of this method, 2) validate the accuracy and precision of perturbation onset timing and duration, and 3) present the stability demands imparted by the method among individuals with impaired balance (i.e. unilateral transtibial limb loss; UTTLL), as quantified by step length, step width, minimum mediolateral margin of stability (ML-MoSmin), and ranges in sagittal whole-body angular momentum (WBAM) in order to relate the destabilizing effect of our perturbation approach to that of other methods. Our rationale for selection of these stability measures is that altering the base of support through altered step lengths and widths is an important, and commonly quantified, strategy used to regulate balance during walking. Further, ML-MoSmin expands on observations of changes to the base of support by incorporating information about center of mass dynamics (16–19). Lastly, sagittal WBAM reflects the ability of an individual to coordinate rotational momentum of the entire body relative to the center of mass, thus providing different but complementary information about the effects of the perturbation on an individual’s dynamics (20).

## 2. Methods

### 2.1 Perturbation Application

The perturbation application was implemented using LabVIEW 2017 (National Instruments, Austin, TX, USA), Bertec Treadmill Control Panel v1.7.3 (Columbus, OH, USA), and a National Instruments USB-6211 DAQ (Figure 1). In the application, analog voltages representing bilateral vertical ground reaction forces (vGRFs) from the signal conditioners of a Bertec treadmill are input to the DAQ and used to estimate heel-strike and toe-off events in real-time. Gait events are detected using a 30 N threshold in vGRFs. These events are used to calculate bilateral stance durations. A running average of the previous five stance durations are stored to inform perturbation timing. When a perturbation is commanded, the stance duration and user-defined perturbation duration, side, and type (i.e. “trip” or “slip”) are used to apply a unilateral belt deceleration (trip) or acceleration (slip). The onset of this perturbation is applied at a user-defined percentage of stance duration following the subsequent heel-strike on the perturbed side. The perturbation application generates an 8-byte unsigned packet representing desired belt velocities and accelerations that are sent to the Bertec Treadmill Control Panel via transmission control protocol (TCP) remote connection. When perturbations are not commanded, the application functions as the standard Bertec Treadmill Control Panel.

**Figure 1:**
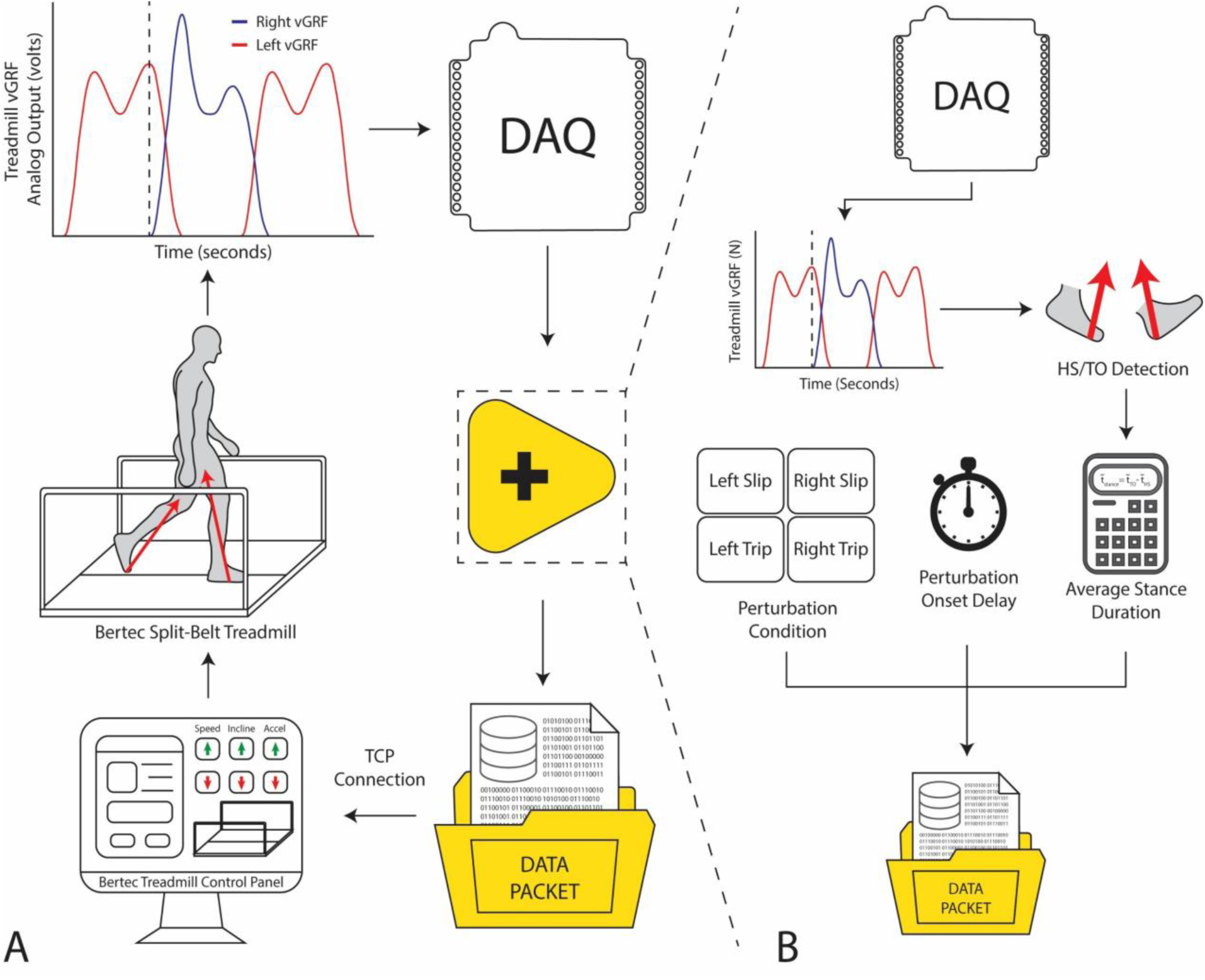
Overview of the closed-loop perturbation application program (A). Vertical ground reaction forces generated during walking on the split-belt treadmill are acquired in real time and streamed to a data acquisition board. These signals are used to detect gait events and trigger perturbation logic, which is packaged into command data and sent to the treadmill control software. The treadmill software then issues velocity and acceleration commands to apply the desired perturbation during walking. (B) Detailed view of the LabVIEW-based control logic. Analog gait signals are used to detect heel-strike (HS) and toe-off (TO) events, from which average stance duration is calculated using the previous five stance durations. User-defined perturbation parameters, including perturbation condition (left/right slip or trip) and onset delay relative to the detected gait event, are used to generate a command data packet. This packet is transmitted via a TCP connection to the treadmill control computer to initiate the desired perturbation.

### 2.2 Experimental Protocol

Five participants with traumatic UTTLL ([mean ± standard deviation] 3M/2F, Age: 38.0 ± 11.2 yrs, Stature: 1.77 ± 0.11 m, Body Mass: 86.1 ± 9.8 kg) walked unperturbed for one minute at 1.0 m/s to acclimate to the treadmill. Following the acclimation period, participants experienced all perturbation conditions (intact-side slip, intact-side trip, prosthetic-side slip, and prosthetic-side trip) while walking at a slower velocity (0.7 m/s). Perturbations consisted of brief (200 ms duration), rapid (10 m/s^2^ acceleration/deceleration) increases (“slips”) or decreases (“trips”) in unilateral treadmill velocity. Target belt velocities for “slips” and “trips” were 2.0 m/s and 0.0 m/s respectively. Desired perturbation onset timing was set to 40% of stance. These perturbation parameters were selected to elicit destabilizing (but not fall inducing) gait disturbances that targeted single stance to allow for comparison of intact and prosthetic limb balance strategies (20–22). Once participants indicated they were comfortable with all perturbations at 0.7 m/s, they repeated the perturbations at 1.0 m/s until they chose to no longer use handrails. Participants then continuously walked for 12-15 minutes with unexpected belt perturbations occurring every 20-35 steps. Each participant experienced a minimum of 40 perturbations (two sides X slip/trip X 10 repetitions) in a random order. Later participants experienced additional perturbations to replace cases of crossover steps or software errors resulting in unexpectedly long perturbations.

Kinematics were collected at 120 Hz using a 22-camera Qualisys motion capture system (Gothenburg, Sweden). Sixty-six markers were used to track full-body motion. Additionally, flat reflective markers were placed ∼20 cm apart along the outside edge of each treadmill belt for velocity validation. vGRFs were collected at 1200 Hz. All participants were secured to a harness to prevent falls during walking.

### 2.3 Validation Metrics

Similar to (20), perturbation onset accuracy/precision was quantified as the mean/standard deviation, respectively, of the elapsed time between the desired perturbation onset time and when measured belt velocity crossed above 1.1 m/s (for slips) or below 0.9 m/s (for trips). The absolute perturbation onset delay was calculated as:

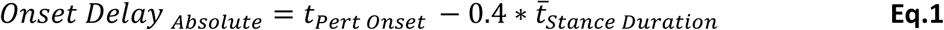

where *t̅_Stance Duration_* is the average stance duration from the previous five steps on the perturbed side prior to the perturbation. *Onset Delay _Relative_* was calculated using:

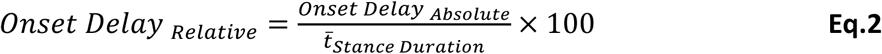

Similarly, perturbation duration accuracy/precision were quantified as the mean/standard deviation, respectively, of the absolute and relative times between perturbation onset and offset. The absolute perturbation duration was calculated as:

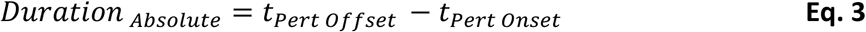

Where *t_Pert Offset_* was the time when belt velocity crossed below 1.1 m/s or above 0.9 m/s while returning to 1.0 m/s for slips or trips, respectively. The relative perturbation duration was calculated as:

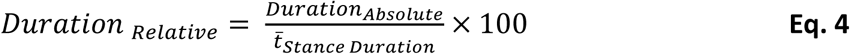

### 2.4 Data Processing

Data were analyzed using MATLAB (Mathworks, Natick, MA USA) and OpenSim v4.4 (SimTK, Stanford, CA, USA). A 23-segment generic full-body musculoskeletal model of an individual with UTTLL (23), revised to include a two-segment prosthetic foot, was scaled to each participant’s anthropometry based on motion capture during a static standing pose. Inertial properties of body segments were scaled by distributing each participant’s measured body mass across body segments such that the mass distribution was preserved relative to the generic model. Marker trajectories and kinetic data were low-pass filtered using a zero-phase 4^th^ order Butterworth filter at 6 Hz and 15 Hz respectively. Gait events were detected using a 30 N threshold applied to vGRFs. Model kinematics were calculated using OpenSim’s Inverse Kinematics and Body Kinematics tools. Step lengths and widths were calculated as the anteroposterior and mediolateral distances, respectively, between the centers of mass of each foot segment at the instance of heel strike.

ML-MoSmin was calculated as the minimum difference between the extrapolated center of mass, XCoM, and the medio-lateral center of pressure, COPML during stance phase (24). XCoM was calculated as:

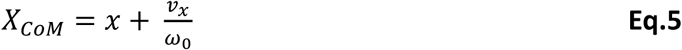

Where *x* is the vertical projection of the center of mass on the ground and *v_x_* is the velocity of the center of mass. *ω*_0_ is the natural frequency of the body modeled as a pendulum:

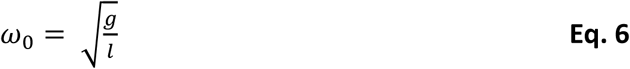

Where *g* is gravitational acceleration and *l* is leg length measured from the anterior superior iliac spine to the midpoint of the lateral and medial malleoli measured during static standing.

WBAM was calculated relative to the whole-body center of mass (COM) using a custom MATLAB script. Segment masses and inertias were obtained from scaled models. WBAM was calculated as (25,26):

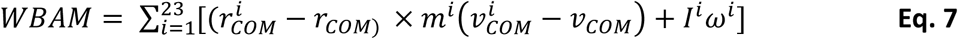

Where *i* indicates the segment number (23 total due to the model), *r_COM_* is whole-body COM position, *r^i^_COM_* is segment COM position, *m^i^* is the mass of the segment, *v^i^_COM_* is the velocity of the segment COM, *v_COM_* is the velocity of the whole-body COM, *I^i^* is the segment inertia tensor, and *ω^i^* is the segment angular velocity. WBAM was expressed in the pelvis reference frame and was normalized by *body mass*walking speed*height*.

### 2.5 Statistics

Paired-sample t-tests assessed mean differences between the perturbed step (“S0”) and the ipsilateral pre-perturbation step (“S-2”) in addition to the recovery step (“S+1”) and the ipsilateral pre-perturbation step (“S-1”) for each perturbation condition (i.e., “slip” or “trip”). Normality was confirmed for all results (absolute skewness < 1.31, absolute kurtosis < 2.97) (27). The critical α was set to 0.05. All paired sample t-tests were performed in MATLAB 2024b.

## 3. Results

### 3.1 Perturbation Validation

Of 221 total perturbations, 26% (58 of 221) were excluded due to: the TCP connection producing perturbations with durations exceeding 150% of the target duration (45 of 221) or participants crossing over belts following a perturbation (13 of 221); resulting in 176 perturbations used for this analysis. For all analyzed trials, the average accuracy and precision of absolute onset delay, measured by the mean and standard deviation, was 183.3 ± 9.7 ms. Relative onset delay of 24.2% ± 1.9% of stance duration. Absolute perturbation duration was 239.9 ± 7.5 ms, 18.1% longer than the target perturbation duration. Relative perturbation duration was 31.6% ± 2.4% of average stance duration. Timeseries plots representing belt velocities are shown in Figure 2.

**Figure 2:**
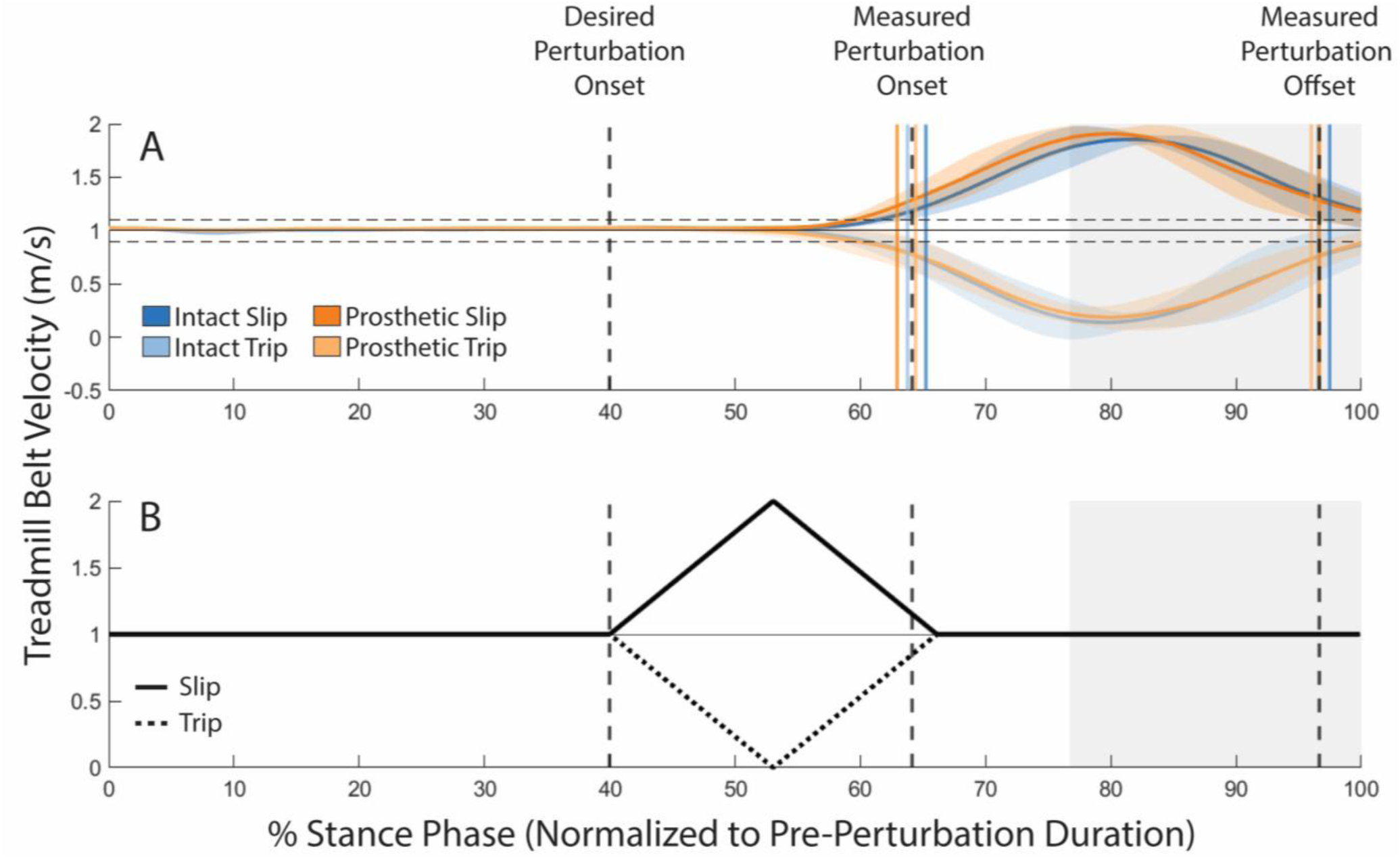
Timeseries plot of (A) measured and (B) theoretical unilateral treadmill belt velocity during each perturbation condition. Black horizontal line indicates unperturbed treadmill belt velocity (1.0 m/s). Dashed horizonal lines indicate 90% and 110% velocity threshold to indicate perturbation onset. Vertical dashed lines indicate desired (40% of stance duration) and measured perturbation onset/offset respectively. Percent stance phase was normalized to pre-perturbation stance duration. Shaded regions indicate ± 1 standard deviation. Grey shaded regions indicate the second double support phase.

### 3.2 Stability outcome measurements

The length of the step immediately following the perturbation (S0) was affected by all perturbation conditions (Fig. 3A). Intact- and prosthetic-side trips decreased perturbed step length by 15.1% (p = 0.005) and 9.4% (p = 0.010) respectively whereas intact- and prosthetic-side slips increased perturbed step length by 7.5% (p = 0.008) and 8.3% (p = 0.005). During the recovery step, step lengths decreased by 35.3% (p = 0.010) and 31.6% (p = 0.008) following intact-side slips and prosthetic-side slips, respectively. Conversely, during the recovery step, step lengths increased by 6.3% (p = 0.017) following prosthetic-side trips. Intact-side trips did not affect the recovery step following a perturbation (p = 0.261).

**Figure 3:**
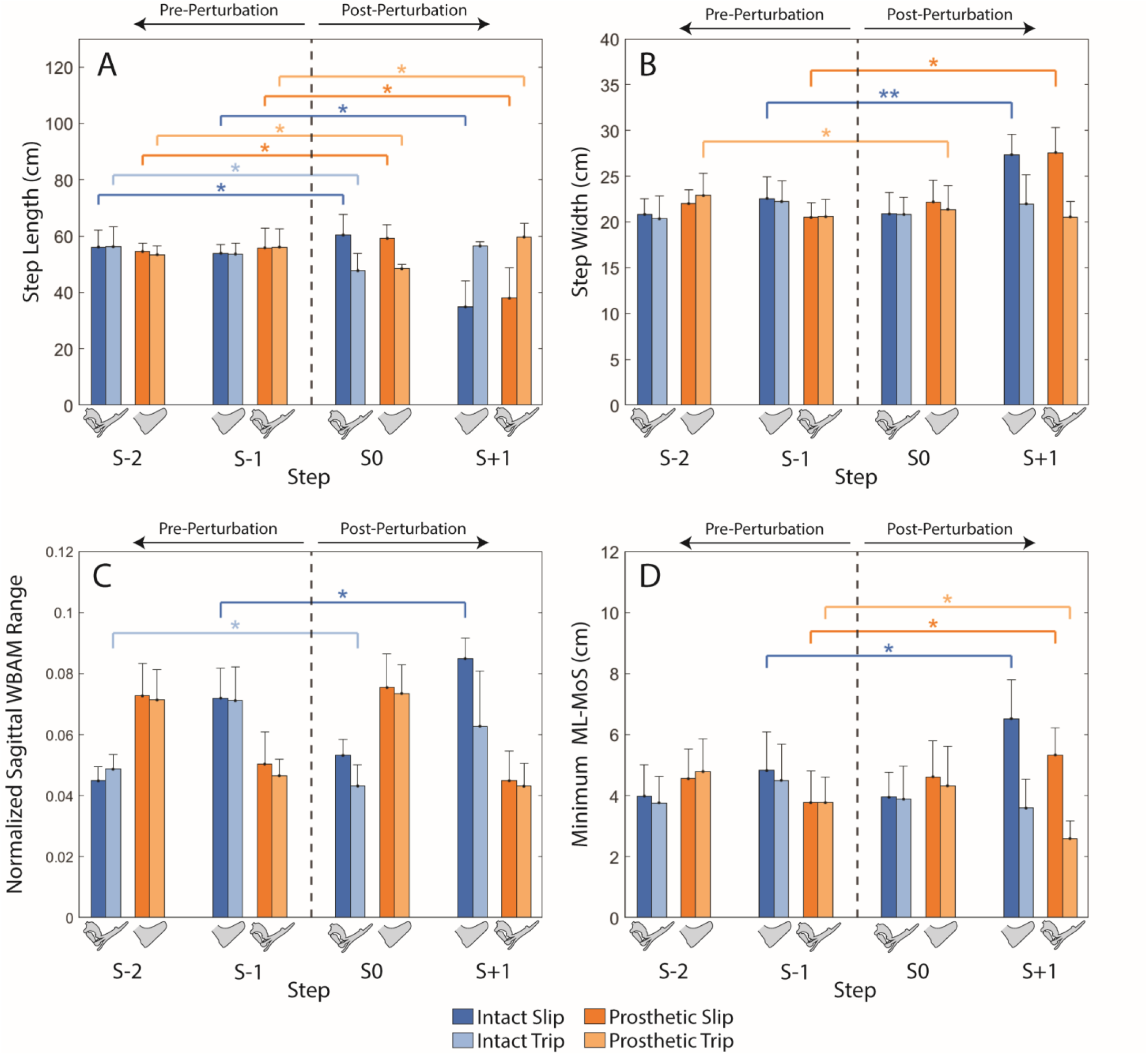
Spatiotemporal and stability measures across steps before and after perturbation. (A) Step length, (B) step width, (C) normalized sagittal whole-body angular momentum (WBAM) range, and (D) minimum mediolateral margin of stability (MoS) are shown for the two steps preceding the perturbation (S–2, S–1), the perturbed step (S0), and the recovery step (S+1). Sagittal WBAM was normalized to *body mass*heigh*velocity*, resulting in an unitless measurement. Perturbations were applied to either the intact or prosthetic limb during slip and trip conditions, as indicated by bar color. S0 corresponds to the step contralateral to the perturbed limb. Error bars denote 1 standard deviation. Horizontal brackets indicate statistically significant pairwise comparisons. * Indicates p < 0.05 and ** indicates p < 0.001

Prosthetic-side trips were the only perturbations to affect step width for the first step after the perturbation (S0; Fig 3B) and were found to decrease step width by 6.6% (p = 0.029). Both the intact- and prosthetic-side slips increased recovery (S+1) step width by 21.2% (p < 0.001) and 34.6% (p = 0.001) respectively. There were no changes in recovery step width resulting from a prosthetic- or intact-side trip (p > 0.883).

Normalized sagittal WBAM range (Fig 3C) decreased by 13.6% (p = 0.003) following intact-side trips (S0). Only the recovery step was affected following an intact-side slip, with sagittal WBAM range increasing by 19.9% (p = 0.023). There were no differences in sagittal WBAM range when a perturbation occurred on the prosthetic-side (p > 0.150), regardless of perturbation type.

ML-MoSmin (Fig 3D) was affected on the recovery step (S+1), increasing by 40.5% (p = 0.003) and 35.2% (p = 0.010) during a prosthetic- and intact side slip respectively, with only the prosthetic-side trip decreasing minimum mediolateral margin of stability 29.4% (p = 0.019). There were no differences during the perturbed step (S0) from either the prosthetic- or intact side (p > 0.052).

## 4. Discussion

The main objective was to present an open-source method to apply perturbations at specified times during stance phase of walking using a standard (Bertec) split-belt treadmill. We described the hardware and software of this method, validated the accuracy and precision of the perturbation onset timing and duration, and quantified the stability demands imparted by the method among individuals with impaired balance (i.e., UTTLL).

To evaluate the method’s accuracy and precision, the desired perturbation onset time and perturbation duration to measured values were compared. On average, the application administered perturbations ∼24% later in stance (183.4 milliseconds) than intended, which is approximately 3 times that of a similar application performed using a high-performance system (CAREN, Motek, Netherlands) (20). The mean absolute perturbation duration was 239.9 ± 12.9 ms, ∼40 ms longer than desired and lasting ∼31.5% of stance phase, ∼5.4% longer relative to the anticipated 26.1% of stance phase. The theoretical treadmill velocity profile and desired onset and duration times demonstrate that the perturbation is expected to terminate prior to the start of double-support (Fig 2B). However, the measured perturbation extended into double-support phase, terminating before the initiation of the swing phase (Fig 2A). These delays are likely due to software and hardware limitations (i.e., sampling rate discrepancy between LabVIEW and the treadmill controller). As the loop rate of the DAQ was approximately 1000 Hz, and the loop rate of the treadmill controller was approximately 50 Hz, future iterations of this perturbation paradigm seeking to reduce delays should focus on optimizing the treadmill controller, in addition to optimizing the TCP connection between the DAQ and Bertec Treadmill Control Panel for higher speed communication.

To assess the effectiveness of the methodology at inducing instability, we quantified the stability demands imparted by the perturbations on individuals with impaired balance. Changes in step length, step width, ML-MoSmin, and sagittal plane WBAM ranges after vs. before perturbations, comparing pre-perturbation steps with post-perturbation steps were assessed. Following a slip, step length increased during the perturbed step, which increased braking force (Fig S1), allowing for greater dissipation of mechanical work at leading limb collision and reducing center of mass velocity (28–32). Due to the reduced center of mass velocity, the recovery step was smaller relative to pre-perturbation steps, providing stability as the participant returns to their steady-state gait (30). The increased step length during the perturbed step and reduced step length during the recovery step is consistent with previous work that has investigated slipping perturbations using belt accelerations among uninjured individuals, ranging between a 1.3% - 12.8% increase during the perturbed step and a 3.1% - 43.2% decrease during the recover step (19–22,33–37).

Conversely, during trips, the opposite effect was observed, with shorter step lengths during the perturbed step and larger step lengths during the recovery step. A shorter step length during the perturbed step may reflect a rapid, stabilizing response aimed at quickly reestablishing balance and preventing a fall (34,36). The increased recovery step length following a trip may serve to address the increased forward pitching of the body relative to the center of mass during trip recovery (Fig S2). However, this was only observed during a prosthetic-side trip. Following an intact-side trip, there was no difference in step length during the recovery step, suggesting that participants rely on alternative recovery mechanisms, such as counter-rotation movement of the trunk, arms, or legs rather than foot placement adjustment (38,39), particularly when the perturbation occurs on their more stable, intact-side.

Step width remained unchanged across most perturbation conditions, with differences observed only after the prosthetic-side trip on the perturbed step, the prosthetic-side slip on the recovery step, and the intact-side slip on the recovery step. This is partially supported by prior work which found 12.7% - 50.8% increases in step width during the recovery step following a slip (19,20,36). However, Shokouhi et al. (2024) reported decreased step width during tripping conditions (36). This discrepancy is likely attributable to methodological differences between studies; specifically, Shokouhi et al. (2024) defined perturbation onset at heel strike, employed slower belt accelerations and decelerations during slip and trip perturbations, and included uninjured young adults.

Sagittal WBAM range was affected only during intact-side slips and trips to an extent similar to that of Shokoushi et al. (2024), who reported a ∼23% and ∼43% increase to WBAM range in uninjured young adults following a slip and trip, respectively (36). However, examination of the time-series of sagittal WBAM reveals influences of the perturbations that were not captured by ranges in sagittal WBAM alone (Fig S2). For trips, negative WBAM (i.e., forward pitching) at the heel strike marking the end of the perturbed step was diminished relative to unperturbed walking, with this arrested forward angular momentum being similar between perturbations delivered on the intact and prosthetic sides. Sagittal external support moments generated by the anteroposterior and vertical GRFs (Fig S5) shows a smaller forward pitching moment at the onset of the trip was due to a smaller propulsive force and larger vGRF of the trailing leg (Fig S1 & Fig S3). However, for slips, negative WBAM at this initial contralateral heel strike was larger for perturbations delivered on the intact side and relatively unchanged from unperturbed levels for perturbations delivered on the prosthetic side. Unexpectedly, this difference in prosthetic-vs. intact-side slip sagittal WBAM was not reflected by prosthetic vs. intact-side differences in step length, suggesting individuals with UTTLL may avoid using their prosthetic foot for braking to arrest increased forward pitching. Such reduced braking is consistent with a general trend of reduced posterior GRFs during unperturbed walking among individuals with vs. without UTTLL (40). During the recovery step following a prosthetic-side perturbation, sagittal WBAM remains negative, indicating continued forward pitching, regardless of the significant change in step-length. This agrees with previous work, reporting a weak correlation between sagittal WBAM and step length following an anterior-posterior perturbation (14,41). However, during intact-side slips, participants seem to “walk through” the perturbation, with the large forward pitching WBAM induced by the perturbation rapidly returning to pre-perturbation values within the recovery step. During the perturbed step following the intact-side slip, participants generate a large 1^st^ peak vGRF (Fig S3) with their prosthetic foot, creating a large positive moment about the CoM (Fig S5C), contributing to the rapid correction of WBAM during early double support-phase. In combination with a quick, shortened but wide recovery step onto their intact leg (Fig 3 & Fig S6), participants are able to rapidly recover from the perturbation using a foot-placement approach, widening base of support, enabling WBAM to quickly return to pre-perturbation values.

ML-MoSmin was largely affected during the recovery step for all perturbation types with the exception of intact-side trips. The increase in ML-MoSmin during the slip-perturbation is consistent with the increased step width observed during the recovery step, creating a larger base of support to provide stability following the perturbation. Hak et al. (2013) observed similar changes to ML-MoSmin applying a medio-lateral perturbation using a CAREN system, observing a ∼10.5% - ∼16.9% increase in ML-MoSmin compared to unperturbed walking in individuals with UTTLL (19). However, following a prosthetic-side trip, we observed a decrease in minimum ML-MoSmin while step width remained unchanged, implying the XCoM approached the intact foot base of support more closely than during unperturbed walking. These differences in ML-MoSmin may depict different “strategies” used by individuals with UTTLL, relying on a “stepping strategy” for foot-placement of the prosthetic leg and an “ankle strategy” to adjust center of mass dynamics using the intact leg (42).

In addition to the aforementioned hardware/software limitations of the perturbation application, there are also several notable limitations to our study design. First, our participants consisted of Service members with UTTLL, due to traumatic injury and thus were generally younger and more active relative to civilians with lower limb loss due to dysvascular etiologies. Therefore, the instability elicited by the perturbation measured in this study may be considered a “lower bound” among balance-impaired populations and is anticipated to be larger among other groups of individuals with UTTLL or balance-impaired populations. Further, due to the high activity level of our participants, we selected a fixed walking speed that was relatively high compared to the self-selected walking speed of “average” individuals with limb loss. Prior reports have identified 0.86 m/s and 1.19 m/s as the average self-selected walking speeds for K3 and K4 ambulators, respectively, based on the Six-Minute Walk Test (43). Finally, all testing and software development was conducted using a single split-belt treadmill, which may limit the generalizability of the findings across other treadmill makes and models.

## 5. Conclusion

In this paper, we introduced an open-source method for delivering perturbations in unilateral belt speed at user-specified times during the gait cycle using a standard (Bertec) split-belt treadmill. This approach successfully induced instability among individuals with impaired balance, demonstrating its feasibility as an open-source alternative to more specialized perturbation approaches. While future work will focus on refinement of the hardware and software to improve perturbation accuracy and consistency, we anticipate the application in its current form will provide the broader research and clinical community with an accessible tool to investigate stability of different populations during walking within safe, highly controlled, and data-rich laboratory environments.

## List of Abbreviations

UTTLL: Unilateral transtibial limb loss
WBAM: Whole body angular momentum
ML-MoSmin: Minimum medio-lateral margin of stability
DAQ: Data acquisition
GRF: Ground reaction force
vGRF: Vertical ground reaction force
COM: Center of mass
XCoM: Extrapolated center of mass
COPML: Medio-lateral center of pressure
TCP: Transmission control protocol

## Declarations

Eligible participants provided written informed consent in person prior to participation in the protocol. The protocol was approved by the Institutional Review Boards of Walter Reed National Military Medical Center (IRB Number: WRNMMC-2024-0460)

## Consent for Publication

Consent for publication of videos has been obtained from participants included in supplementary videos.

## Availability of data and materials

The datasets generated and/or analyzed during the current study are available at https://doi.org/10.5281/zenodo.19825271. The perturbation application LabVIEW VI is available at https://github.com/kiash24/Bertec_LabVIEW_SlipTrip

## Author Contribution

CMB conceptualized the study, acquired funding, administered the project, and supervised resources, PRG conceptualized the study and designed the experimental protocol; CMB, PRG, and KFA carried out the experiments; KFA developed the perturbation application, analyzed study data, and drafted the manuscript. CMB, PRG, KFA, and BDH reviewed and edited the manuscript. All authors read and approved the final manuscript.

## Acknowledgements

The authors would like to thank Krista Clark, Julian Acasio, and Paige Agnew for their assistance with data collection. The views expressed herein are those of the authors and do not reflect the views of Henry M. Jackson Foundation for the Advancement of Military Medicine, Inc., Uniformed Services University of the Health Sciences, Defense Health Agency, Department of War, nor the U.S. Government. The identification of specific products or scientific instrumentation is considered an integral part of the scientific endeavor and does not constitute endorsement or implied endorsement on the part of the authors, Department of War, or any component agency.

