## Supplementary Material for "An open-source application for applying rapid transient perturbations using a split-belt treadmill"

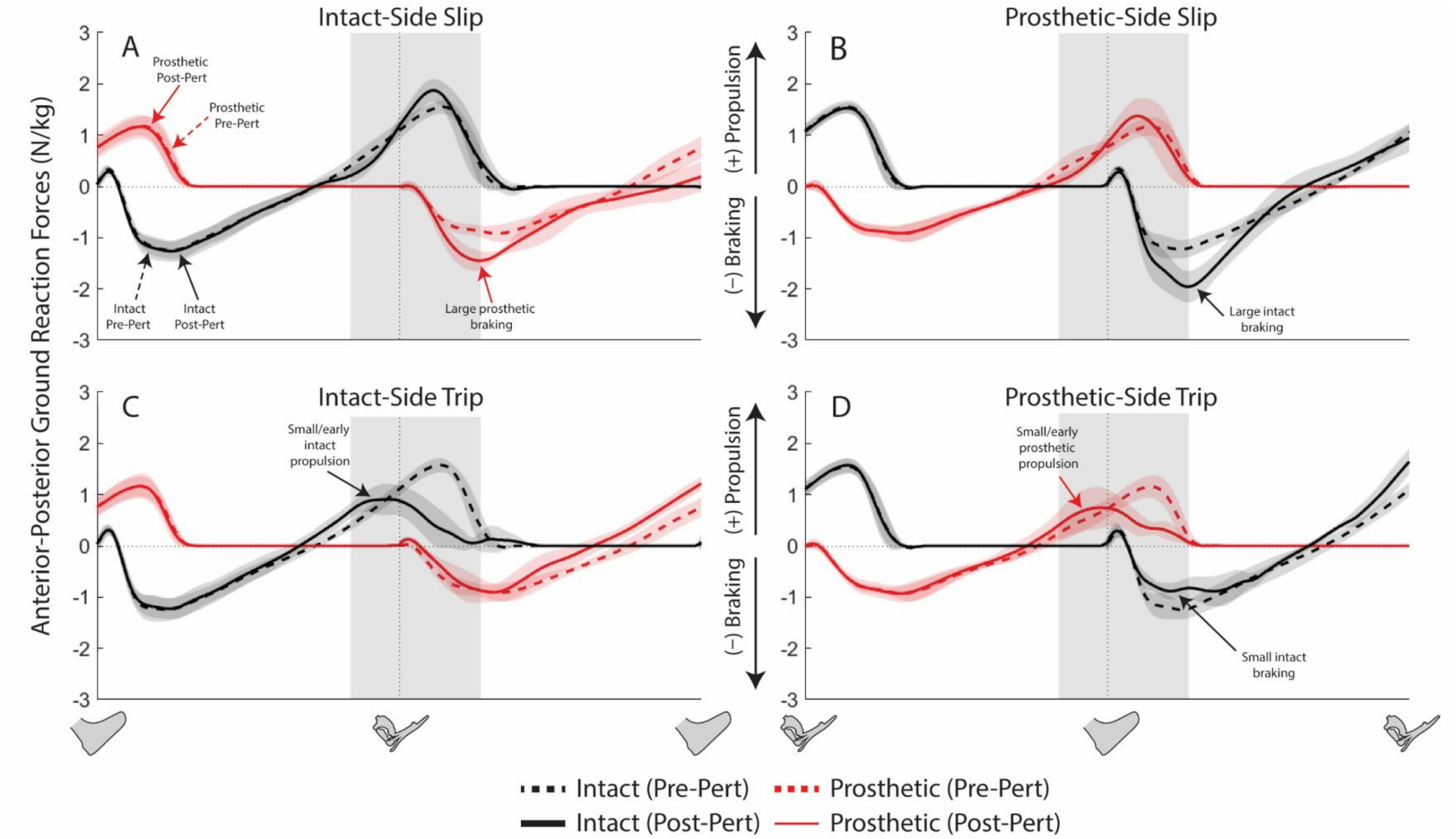


**Figure S1**: Timeseries plot of pre-perturbation anterior-posterior ground reaction forces (AP-GRFs) overlayed on top of post-perturbation AP-GRFs during an (A) intact-side slip, (B) prosthetic-side slip, (C) intact-side trip, and (D) prosthetic-side trip. AP-GRF is normalized to body mass. Dashed lines represent pre-perturbation AP-GRFs. Solid-lines represents the perturbation step and recovery step AP-GRFs. Shaded region around lines indicate ± 1 standard deviations. Vertical grey-shaded region indicates average perturbation onset/offset. Vertical dashed-lines and prosthetic/intact foot indicate heel-strikes.

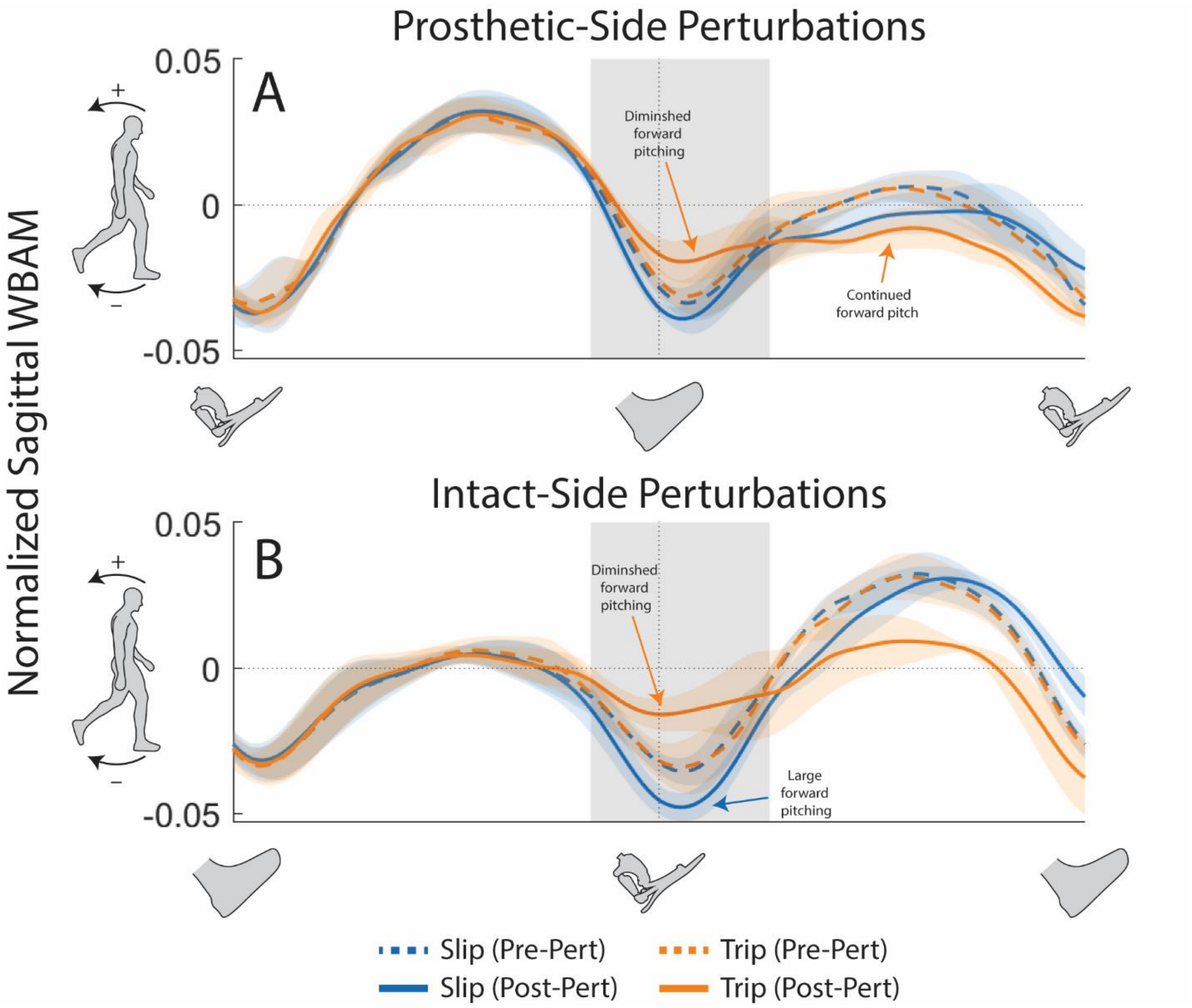


**Figure S2**: Timeseries plot of (A) treadmill belt velocity (m/s) and normalized sagittal WBAM during (B) prosthetic-side and (C) intact-side perturbations. Sagittal WBAM was normalized to *body mass*height*velocity*. Dashed lines and solid lines indicate pre- and post-perturbation values, respectively. Color represents perturbation type (slip/trip). Color-shaded regions indicate ±1 standard deviation. Gray-shaded region indicates average perturbation onset and duration. Vertical dashed-lines and prosthetic/intact foot cartoons indicate heel-strikes.

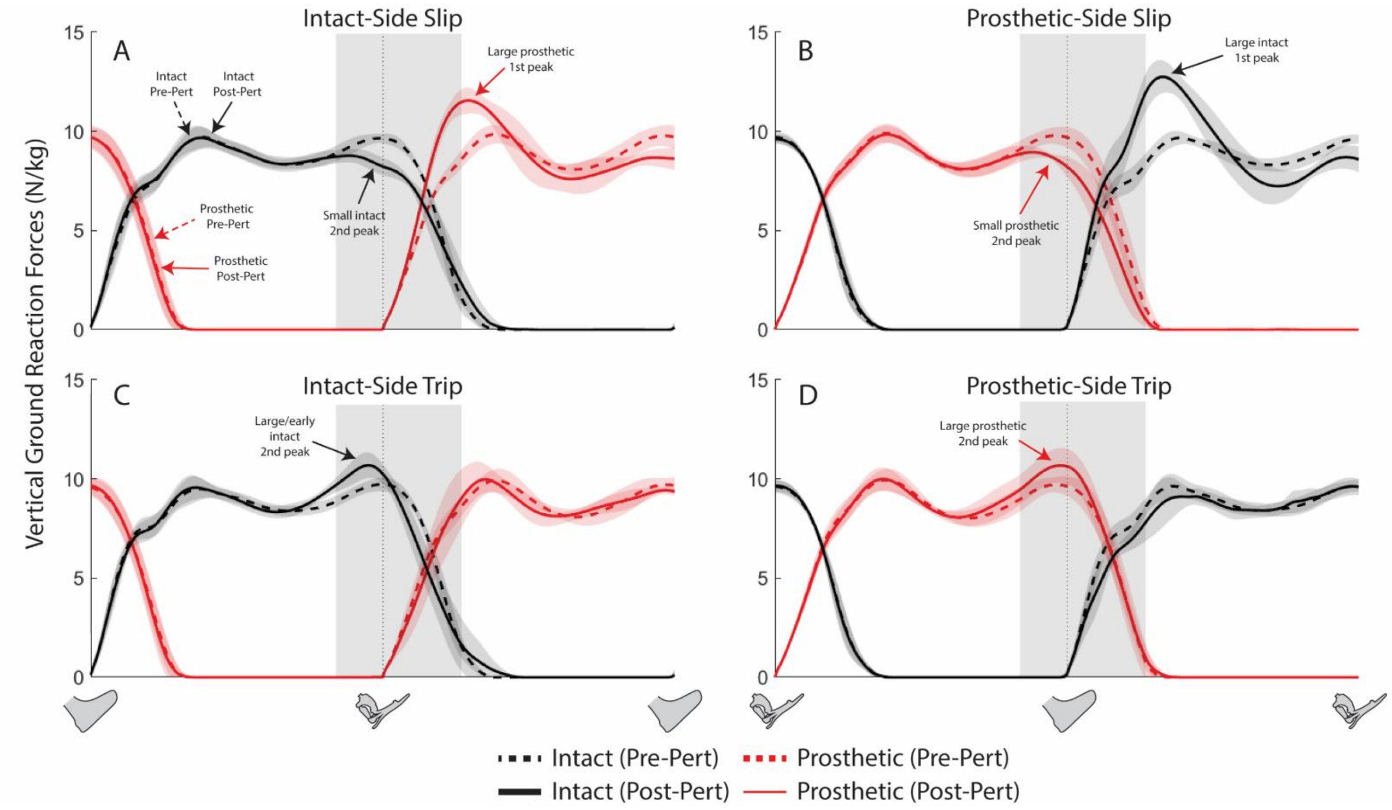


**Figure S3**: Timeseries plot of vertical ground reaction forces (vGRF) during an (A) intact-side slip, (B) prosthetic-side slip, (C) intact-side trip, and (D) prosthetic-side trip. vGRF is normalized to body mass. Solid line represents pre-perturbation (S-2,S-1) vGRF. Dashed-curve represents post-perturbation(S0,S+1) vGRF. Shaded region around lines indicate ± 1 standard deviations. Vertical grey-shaded region indicates average perturbation onset/offset. Vertical dashed-lines and prosthetic/intact foot indicate heel-strikes.

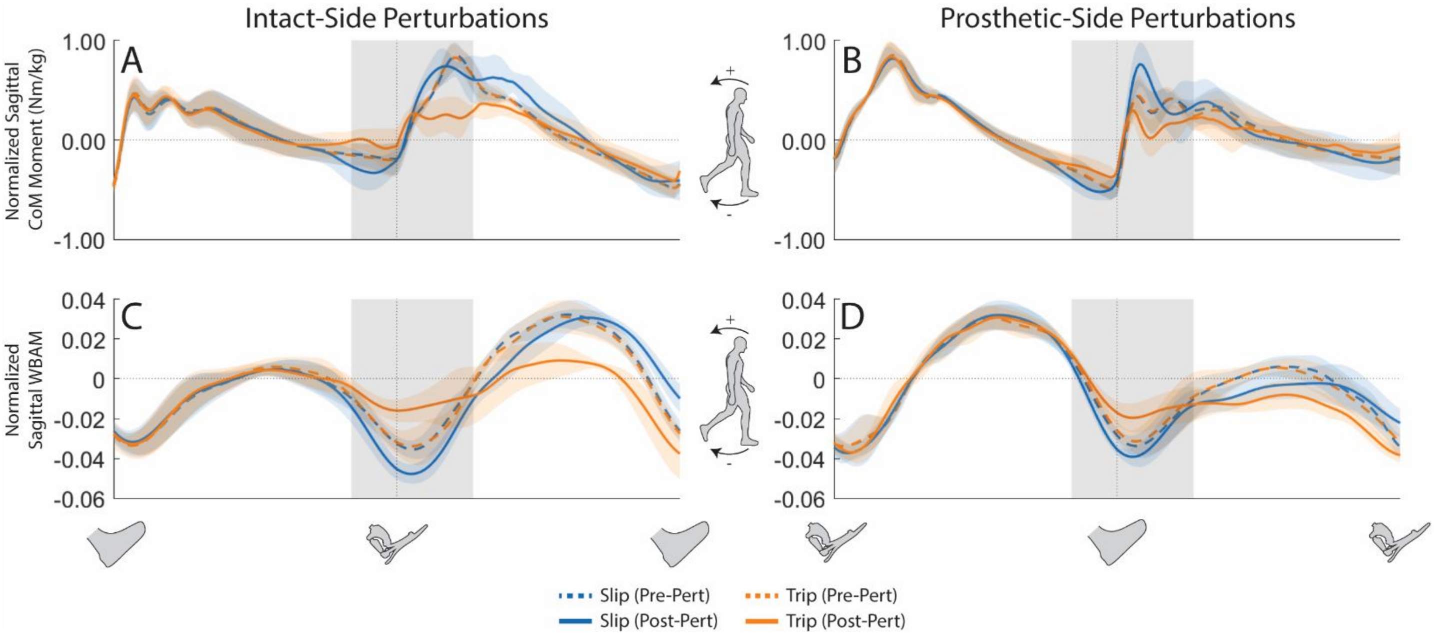


**Figure S4**: Comparison of (A/B) normalized sagittal center of mass moment perturbations versus (C/D) normalized sagittal whole-body angular moment during intact-side and prosthetic-side perturbations. Positive CoM moment/WBAM indicates backward-pitch and a negative CoM moment/WBAM indicates a forward-pitch. Shaded gray-region indicates average time and duration of perturbation. Shaded colored-region indicates ± 1 standard deviation. Dashed-lines represent pre-perturbation steps. Solid-lines represent post-perturbation steps.

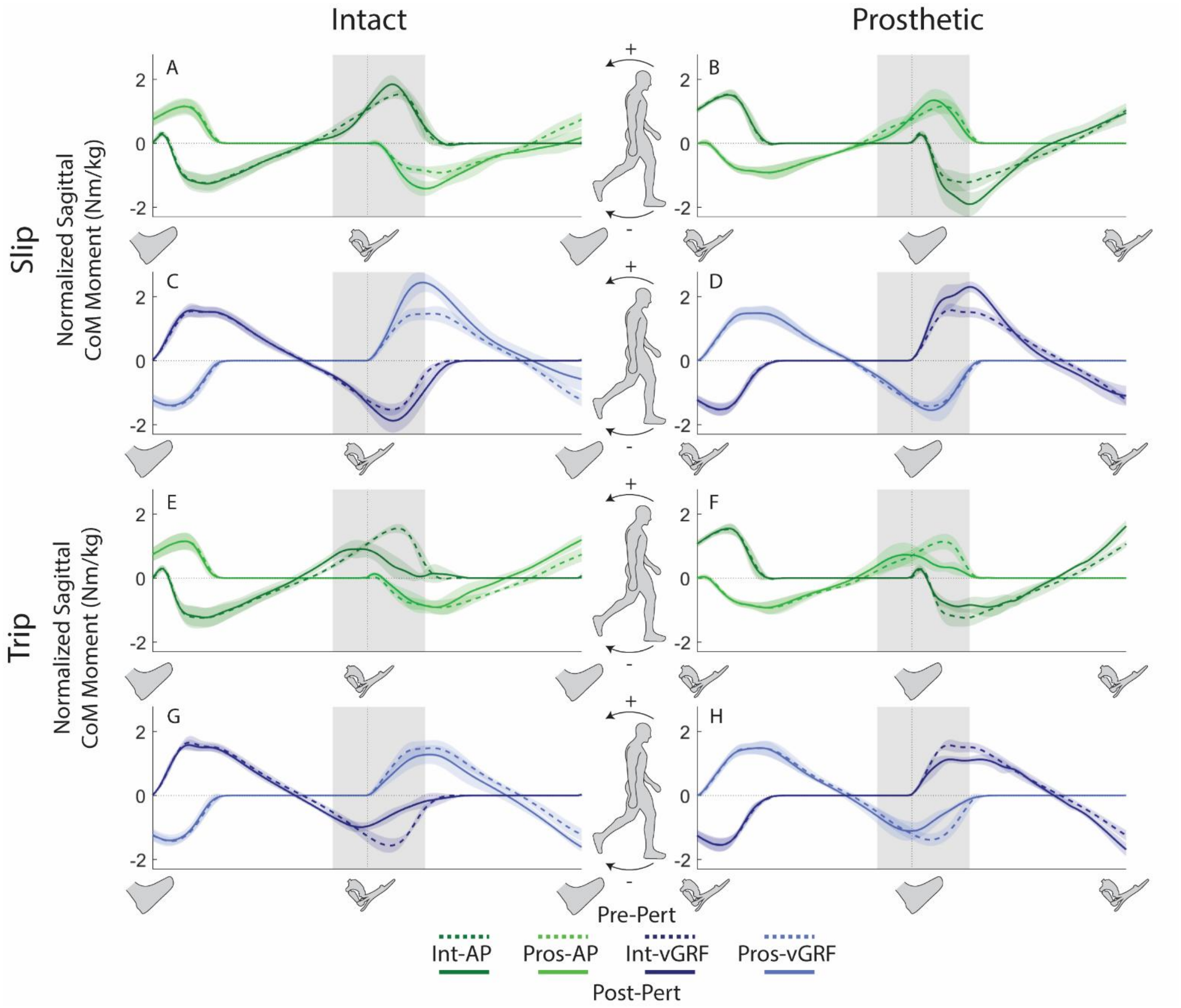


**Figure 1**: Components of the sagittal center of mass momentum during (A/C) intact-slips, (B/D) prosthetic-slips, (E/G) intact-trips, and (F/H) prosthetic-trips. Dashed-line and solid-lines represent pre-perturbation and post-perturbation steps respectively. Gray-shaded region indicates average onset and duration of a perturbation. Color-shaded region indicates $\pm 1$ standard deviation. Positive moment indicates backward-pitch. Negative moment indicates forward-pitch. Heel-strikes are indicated with prosthetic and intact feet.

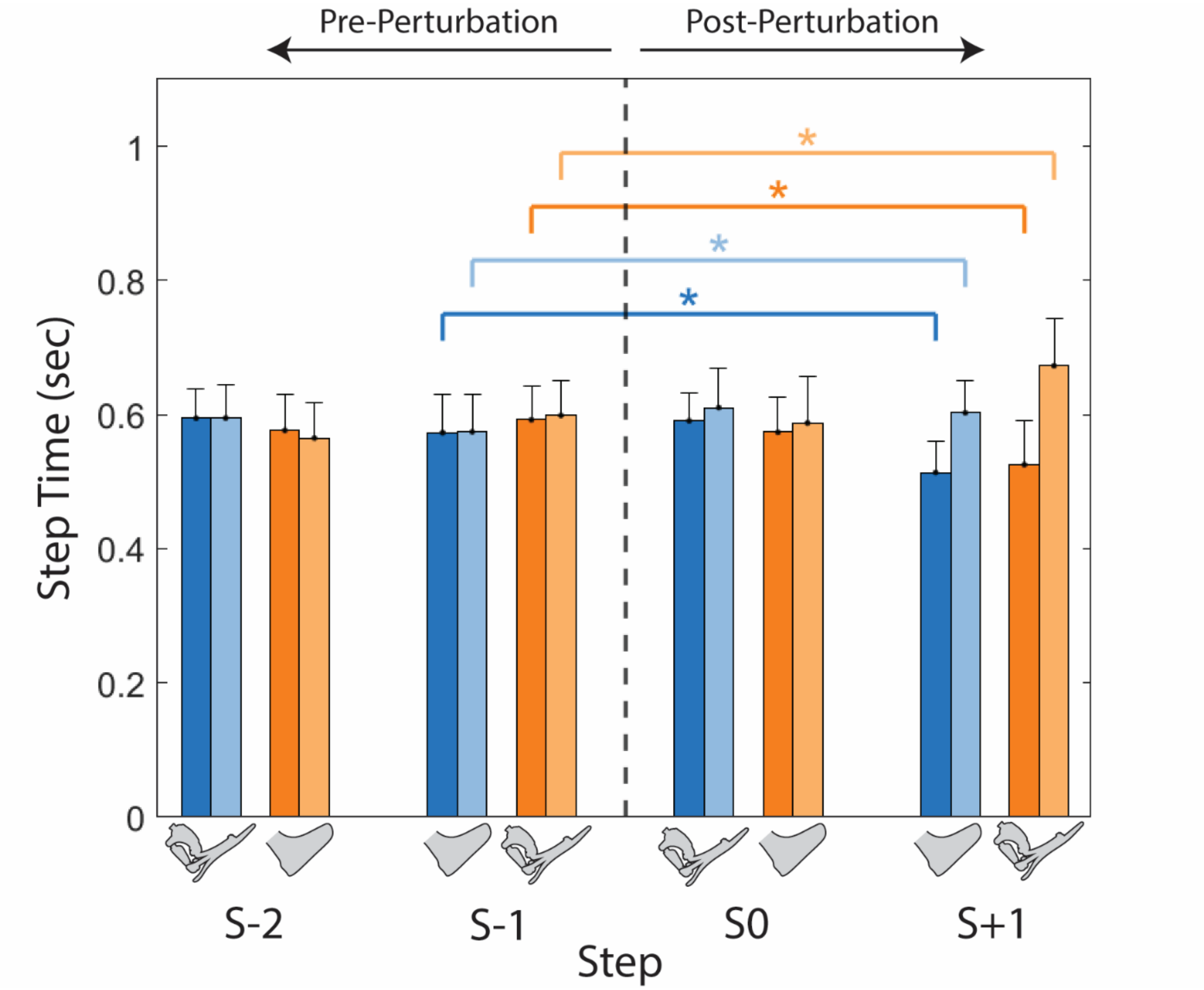


**Figure S6**: Step times before and after perturbation are shown for the two steps preceding the perturbation (S–2, S–1), the perturbed step (S0), and the recovery step (S+1). Perturbations were applied to either the intact or prosthetic limb during slip and trip conditions, as indicated by bar color. S0 corresponds to the step contralateral to the perturbed limb. Error bars denote 1 standard deviation. Horizontal brackets indicate statistically significant pairwise comparisons. * Indicates $p < 0.05$ and ** indicates $p < 0.001$
